# The Effect of Aging, Parkinson’s Disease, and Exogenous Dopamine on the Neural Response Associated with Auditory Regularity Processing

**DOI:** 10.1101/647628

**Authors:** Abdullah Al Jaja, Jessica A. Grahn, Björn Herrmann, Penny A. MacDonald

**Author notes:** Co-senior authors with equal contribution (Björn Herrmann, Penny A. MacDonald). **Correspondence to:** Abdullah Al Jaja, Brain and Mind Institute, Western Interdisciplinary Research Building, Room 5166, University of Western Ontario, London, Ontario, Canada, N6A 5B7. **Abbreviations:** L-3,4-dihydroxyphenylalanine (L-dopa), Parkinson’s disease (PD), Ventral tegmental area (VTA).

## Abstract

Processing regular patterns in auditory scenes is important for navigating complex environments. Electroencephalography (EEG) studies find enhancement of sustained brain activity, correlating with the emergence of a regular pattern in sounds. How aging, aging-related diseases such as Parkinson’s disease (PD), and treatment of PD affect this fundamental function remain unknown. We addressed this knowledge gap. Healthy younger and older adults, and PD patients listened to sounds that contained or were devoid of regular patterns. Healthy older adults and PD patients were tested twice—on and off dopaminergic medication in counterbalanced order. Regularity-evoked, sustained EEG activity was reduced in older, compared to younger adults. PD patients and older controls had comparable attenuation of the sustained response. Dopaminergic therapy further weakened the sustained response in both groups. These findings suggest that fundamental regularity processing is impacted by aging-related neural changes but not those underlying PD. The finding that dopaminergic therapy attenuates rather than improves the sustained response coheres with the dopamine overdose response and implicates brain regions receiving dopamine from the ventral tegmental area in regularity processing.

## 1. Introduction

Auditory scene analysis is the ability to perceptually organize superimposed sounds (i.e., a sound mixture) into meaningful streams, such as a person’s voice or an approaching car (Bregman, 1990). Regular patterns in sounds support perceptual organization as they enable a listener to a) track a sound over time while relegating to the background non-patterned, irrelevant sounds (Griffiths et al., 1998; Winkler et al., 2009), b) predict future events (Barlow, 2001; Bendixen et al., 2009; Denham and Winkler, 2006; Sloutsky, 2003; Sohoglu and Chait, 2016; Turk-Browne et al., 2009), and c) efficiently organize the auditory scenes (Andreou et al., 2011; Bendixen et al., 2007; Christiansen, 2014).

One line of research that focuses on the processing of regular patterns showed that when listeners are exposed to sounds a) containing a regular frequency modulation (Bjorn Herrmann and Johnsrude, 2018), b) made of chords (Teki et al., 2016), or c) made of repeating tones (Sohoglu and Chait, 2016; Southwell et al., 2017), a low-frequency sustained response is elicited in electroencephalography (EEG) or magnetoencephalography (MEG) recordings, relative to the neural response to a sound without a regular pattern (i.e., a random sequence). Further, using MEG and functional magnetic resonance imaging (fMRI), regular relative to random patterns correlate with preferential brain activation in hippocampus, superior temporal gyrus, and inferior frontal gyrus (Barascud et al., 2016). Therefore, the processing of regular patterns in sounds appears to rely on a variety of cortical structures, including auditory cortex (Lütkenhöner et al., 2011) as well as brain regions supporting higher-level functions, such as the inferior frontal gyrus and hippocampus (Barascud et al., 2016).

Aging and neurodegenerative disorders are speculated to interfere with regularity processing. Previous investigations of auditory scene analysis in aging have revealed difficulty for the aging population to temporally segregate simultaneous auditory stimuli (Alain et al., 1996) as well as difficulty in inhibiting irrelevant auditory information (Henry et al., 2017). In addition, it has been suggested that aging might affect speech perception and comprehension due to impairment in auditory processing resulting in the reallocation of mental resources towards identifying words, compromising the understanding and evocation of meaning of that word (Pichora-Fuller, 2003). Other studies have identified deficits, such as loss of temporal precision (Anderson et al., 2012), or of automatic discrimination (Getzmann and Näätänen, 2015) in the auditory scene analysis. These studies demonstrate a deficit in temporal processing that occurs during aging, suggesting a loss of temporal precision (Anderson et al., 2012). These studies support a deficincy in temporal processing related to detecting regularities in the aging population.

Interestingly, patients with Parkinson’s disease (PD) are also impaired in temporally segregating auditory stimuli and have deficiency in auditory-to-motor entrainement (Meijer et al., 2016; te Woerd et al., 2018). Though there are fewer studies, these findings suggest that PD might be associated with a deficiency in regularity processing within scenes.

PD is a common, debilitating neurodegenerative disease that principally affects the basal ganglia, a collection of sub-cortical nuclei that control movement and aspects of cognition (Dauer and Przedborski, 2003; Hirsch et al., 2016). In PD, there is profound loss of dopamine-producing neurons in the substantia nigra pars compacta (SNc). The SNc provides dopamine to the dorsal aspect of the striatum (DS, i.e., the bulk of the caudate nucleus and putamen). Dopamine deficiency to the DS produces the cardinal motor manifestations of PD including bradykinesia (i.e., slowness of movement), resting tremor, and muscular rigidity (Dauer and Przedborski, 2003), as well as cognitive deficits in cognitive control such as impairments in attentional shifting, response inhibition, and task switching (Macdonald and Monchi, 2011; Solís-Vivanco et al., 2011). Consistent with the presumed pathophysiological basis for these motor and cognitive deficits, L-3,4-dihydroxyphenylalanine (L-dopa), a dopamine precursor that crosses the blood-brain barrier, and dopamine agonist medications, improve these motor (Birkmayer and Hornykiewicz, 1961; Cotzias et al., 1969) and cognitive functions (Hiebert et al., 2019; Macdonald and Monchi, 2011) in PD.

In contrast, not all functions are improved by dopaminergic therapy. In fact, functions such as memory encoding (MacDonald et al., 2013), association learning (Hiebert et al., 2014; Vo et al., 2014), reward processing (Cools, 2006; Macdonald et al., 2013; Macdonald and Monchi, 2011; Vo et al., 2014) are intact at baseline in PD relative to healthy older controls. These cognitive functions are actually worsened by dopaminergic therapy in PD (Cools, 2006; Gotham et al., 1984; Hiebert et al., 2019; Kish et al., 1988; Macdonald et al., 2013; Swainson et al., 2000; Yang et al., 2018), as well as in healthy elderly and young controls (Fungwe et al., 1992; Olefins et al., 1986; Vo et al., 2016). The *dopamine overdose hypothesis* is a prevaling explanation for this effect of dopaminergic therapy in PD. In contrast to the degeneration of the SNc, dopamine-producing neurons in the ventral tegmental area (VTA) are relatively spared in PD (Kish et al., 1988). The VTA provides essentially normal levels of dopamine to the ventral striatum (VS), composed of the nucleus accumbens ventral caudate and putamen, as well as to the limbic (e.g., hippocampus) and prefrontal cortices. According to the dopamine overdose hypothesis, dopaminergic therapy, titrated to motor symptoms and hence DS dopamine deficiency, distributes in a non-targeted fashion, presumably overdosing VTA-innervated brain regions and worsening functions performed by these brain regions (Cools, 2006; Gotham et al., 1984; Kish et al., 1988; Macdonald and Monchi, 2011) The dopaminergic system has been implicated in processing regularities in sounds through its downstream projections, including areas of the prefrontal cortex (PFC) and hippocampus (Barascud et al., 2016; D’Ardenne et al., 2012; Li et al., 2018). Extrapolations from these studies notwithstanding, the effect of aging, PD, and dopaminergic medication on the fundamental process of regularity detection in sound is not well understood.

The aim of this study was to investigate the effect of aging, PD, and exogenous dopamine (e.g., L-dopa) on regularity processing. We measured EEG-evoked responses for elderly controls and PD patients while they listened to tone sequences that either contained a regular auditory pattern or were entirely random. Both Healthy elderly control participants and PD patients performed this EEG-auditory processing task both off and on exogenous dopamine therapy. Previously-collected data in healthy younger controls, in the OFF dopaminergic therapy state only (Bjorn Herrmann and Johnsrude, 2018), were compared to our data to investigate the effect of aging. If aging impacts regularity processing in auditory scene analysis, we expect EEG signal differences in our healthy elderly controls compared to healthy younger participants in the OFF state. Moreover, if regularity processing depends upon DS and/or DS’s cortical partners, regularity processing is expected to be a) impaired in PD patients at baseline, in the OFF state, relative to age-matched controls, and b) improved, even potentially normalized in PD patients tested in the ON state, relative to elderly control participants’ performance. In contrast, if regularity processing depends upon VTA-innervated brain regions such as VS, limbic, or prefrontal cortices, regularity processing is predicted to be comparable in PD and elderly controls off dopaminergic therapy and worsened by the administration of dopamine, due to dopamine overdose, in both groups.

## 2. Materials and methods

### 2.1. Participants

Nineteen PD patients, (7 females; SEM = 66.5±6.1 years), 19 age- and education-matched healthy controls (aCT; 12 females, 65.7±6.7 years) were recruited for this study. Data from one PD patient was excluded due to the use of a hearing aid and data from two healthy controls were excluded because they did not complete both sessions. Health and demographic information for aCT and PD patients are presented in Table 1. Data from 16 healthy young controls (yCT; 10 females, 18.4 ± 0.9 years) collected in a previous study using an identical procedure (Bjorn Herrmann and Johnsrude, 2018). were included in our OFF-state analyses.

**Table 1:** Demographic, clinical information, and cognitive and affective measures for aCT and participants with PD.

|  |  | Group |  | *p* value |
| --- | --- | --- | --- | --- |
|  |  | aCT | PD |  |
| N |  | 18 | 18 |  |
| Age |  | 66.89 (6.25) | 65.94 (5.72) | 0.64 |
| Disease Duration |  | N/A | 6.78 (4.89) |  |
| L-dopa |  | N/A | 311.85 (227.94) |  |
| MOCA |  | 28.33 (1.60) | 27.61 (1.81) | 0.21 |
| ANART |  | 124.40 (8.55) | 123.80 (5.48) | 0.8 |
| Oxford happiness scale |  | 4.00 (1.627) | 3.42 (0.68) | 0.22 |
| BAI | OFF | 1.94 (2.58) | 10.27 (5.49) | <0.01* |
|  | ON | 1.61 (2.06) | 8.00 (4.91) | <0.01* |
| BDI-II | OFF | 4.44 (4.40) | 10.11 (5.81) | <0.01* |
|  | ON | 4.61 (4.27) | 10.50 (6.51) | <0.01* |
| Apathy | OFF | 9.38 (4.43) | 12.00 (5.89) | 0.12 |
|  | ON | 8.88 (3.61) | 13.17 (6.53) | 0.02* |
| UPDRS | OFF | 1.18 (0.42) | 26.91 (1.22) | 0.168 |
|  | ON | 0.96 (0.31) | 22.26 (1.19) | <0.01* |

Data are presented as mean (SEM). MOCA: Montreal Cognitive Assessment; ANART: National Adult Reading Test; BAI: Beck Anxiety Inventory; BDI-II: Beck Depression Inventory-II; Apathy: Starkstein Apathy Scale. UPDRS: Unified Parkinson's Disease Rating Scale. aCT = age matched healthy Controls, PD = Parkinson's disease patients

All participants with PD were previously diagnosed by a licensed neurologist, had no coexisting diagnosis of dementia, or another neurological or psychiatric disease, and met the core assessment for surgical interventional therapy and the UK Brain Bank criteria for the diagnosis of idiopathic PD (Hughes et al., 1992). All PD patients were treated with regular dopaminergic therapy. ACT were within five years of age (average difference was 3.6 years) and three years of education (average difference was 2.4 years) of their matched PD patient (*p* = 0.653 and *p* = 0.342, respectively). No controls were treated with any dopamine-modulating therapy. Participants with PD were recruited through the Movement Disorders Database at the London Health Sciences Centre, Ontario, Canada. Participants abusing alcohol, prescription or illicit drugs, or taking cognitive-enhancing medications including donepezil, galantamine, rivastigmine, memantine, or methylphenidate, or who presented any hearing impairment were excluded from participating in this study.

The motor sub-scale of the Unified Parkinson’s Disease Rating Scale (UPDRS) was scored by a licensed neurologist with sub-specialty training in movement disorders (P.A.M.) to assess the presence and severity of motor symptoms for all patients both off and on dopaminergic medication. ACT were also screened with UPDRS motor sub-score to rule out undiagnosed neurological illness. Mean group demographic, as well as cognitive and affective screening scores for all patients and controls in each experimental group are presented in Table 1. UPDRS motor subscale scores off and on dopaminergic therapy, daily doses of dopamine replacement therapy in terms of L-dopa equivalents (LED), and mean duration of PD were also recorded and presented in Table 1. Calculation of daily LED for each patient was based on the theoretical equivalence to L-dopa(mg) as follows: L-dopa dose(mg) × 1 + L-dopa controlled release(mg) × 0.75 + L-dopa(mg) × 0.33 if on entacapone(mg) + amantadine(mg) × 0.5 + bromocriptine(mg) × 10 + cabergoline(mg) × 50 + pergolide(mg) × 100 + pramipexole(mg) × 67 + rasagiline(mg) × 100 + ropinirole(mg) × 16.67 + selegiline(mg) × 10 (Wullner et al., 2010).

All participants provided written informed consent to the protocol before beginning the experiment. The study was conducted in accordance with the Declaration of Helsinki (World Medical Association, 2013). This study was approved by the Health Sciences Research Ethics Board of the University of Western Ontario.

### 2.2. Protocol

ACT and PD patients were recruited for two recording sessions, conducted at the Brain and Mind Institute at the University of Western Ontario. During the OFF session, PD patients were instructed to abstain from taking their dopaminergic medication as follows: L-dopa/carbidopa and entacapone were withheld for 12-18 hours and dopamine agonists such as pramipexole, ropinirole, or pergolide, as well as amantadine, rasagiline, and selegiline were not taken for 16–24 hours before testing. Healthy controls took a placebo consisting of cornstarch in the OFF state. During the ON Session (ON), patients were instructed to take their medications as prescribed by their treating neurologist. In the ON Session, healthy controls took 100 mg of L-dopa and 25 mg of carbidopa at the beginning of the session, followed by a waiting period of 45 minutes to maximize medication absorption and peak plasma dopamine levels for experimental testing. Placebo and L-dopa were given to aCT in identical capsules prepared by a lab member who was not involved in the study to ensure blinding of both the participant and the experimenter. The OFF and ON sessions were counterbalanced across all participants. The OFF-ON order of the aCT corresponded to the OFF-ON order of the PD patient to whom s/he was matched. ACT were instructed to abstain from consuming heavy meals, alcohol, and/or caffeine the night before and the morning of each session, so there was minimal interference with dopamine absorption. Patients and healthy elderly controls completed questionnaires assessing anxiety, depression, and apathy. Cognitive function was also assessed using the Montreal cognitive assessment (MoCA), American National Adult Reading Test (ANART). At the end of the waiting period (45 minutes) during both sessions, all participants performed the UPDRS Motor Sub-scale. Physiological measures, such as the heart rate and blood pressure were collected twice—once at the beginning and once at the end of each session. At the beginning of each session, a participant’s hearing ability threshold was estimated for each ear using pure-tone audiometry (Fig. 1). YCT were not tested on dopamine treatment, as the data were collected in a previous study (Bjorn Herrmann and Johnsrude, 2018). YCT participants selfreported normal-hearing abilities but these were not explicitly testing given the low likelihood of hearing impairment at the young mean age of this group.

**Figure 1:**
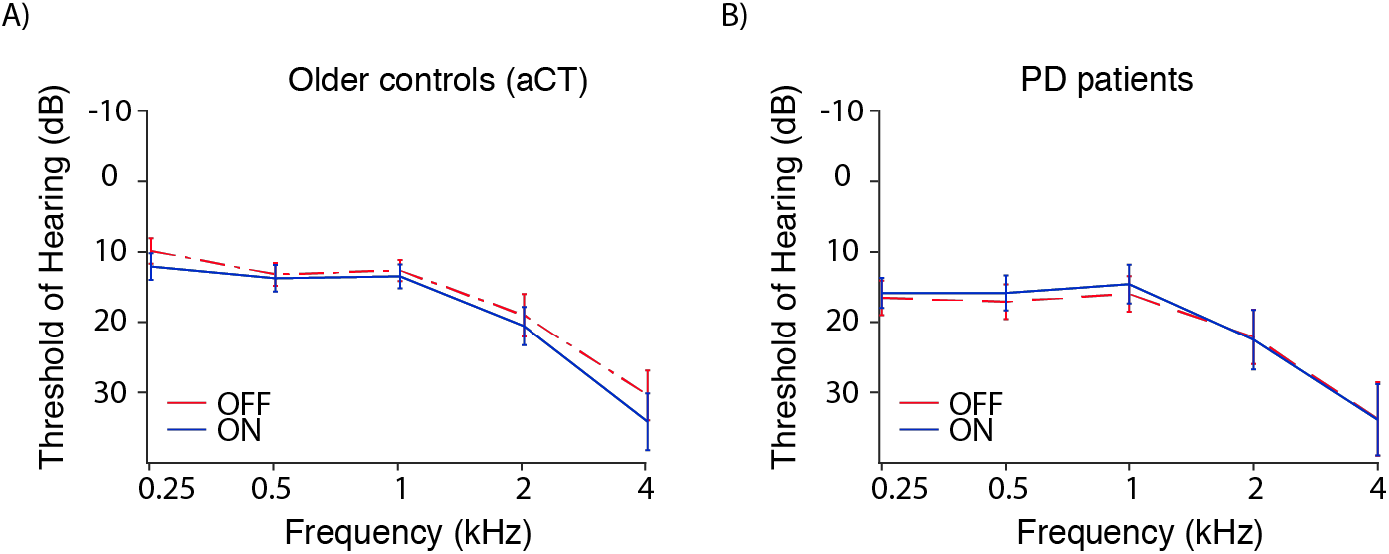
Audiometric data for aCT and PD patients and OFF and ON medication. (A) Mean audiograms for aCT patients OFF (*dotted red line*) and ON (*solid blue line*) are presented, collapsed across left and right ears. (B) Mean audiograms for PD patients OFF (*dotted red line*) and ON (*solid blue line*) medication are presented, collapsed across left and right ears. There were no significant differences between groups or related to dopaminergic therapy in the hearing threshold across the frequencies of our tone stimuli in this study. Error bars represent standard error about the mean (SEM).

### 2.3. Acoustic stimulation

Participants watched a muted movie with subtitles while they were passively presented with sounds via Sennheiser HD25SP II headphones and a Steinberg UR22 (Steinberg Media Technologies) external sound card. Stimulus presentation was controlled by a PC (Windows 7, 64 bit) running Psychtoolbox in MATLAB (R2015b).

Acoustic stimuli were similar to those used in previous studies (Barascud et al., 2016; Björn Herrmann and Johnsrude, 2018; Southwell et al., 2017). Stimuli were 4-s-long sequences made of 0.04-s tones (0.007 s rise time; 0.007 s fall time; 100 tones per sound; no gap between tones). Each stimulus sequence comprised 10 sets of tones and each set was 10 tones (0.4 seconds) long. The frequency of a tone could take on one of 70 values ranging from 700 to 2500 Hz (logarithmically spaced).

The selection of frequency values for each of the 10 sets depended on the stimulus condition. Two conditions were created. For the RAND condition, containing no regularity, 10 new frequency values were randomly selected for each of the 10 sets (Fig. 2, Panel A). In the REG condition, 10 new frequency values were randomly selected for each of the first 4 sets (0–1.6 seconds identical to RAND condition), and then 10 new random frequency values were selected and repeated for set 5 through10, thereby creating a regularity. That is, the serial order of tone frequencies was identical in the repeating sets between 5 and 10 (Fig. 2, Panel B). In this way, in the REG condition, the tones transformed from a random to a regular sequence of tones. Selection of these 10 frequency values and their serial order, used to create the regularity, differed from trial to trial. Each stimulus was presented 108 times over the duration of 3 blocks and individual stimuli were separated by an inter-stimulus interval of 1.5 s. Trials for each condition were presented in a randomized fashion within each block. Stimuli from the RAND and REG conditions were presented along with other auditory stimuli that will not be discussed here.

**Figure 2:**
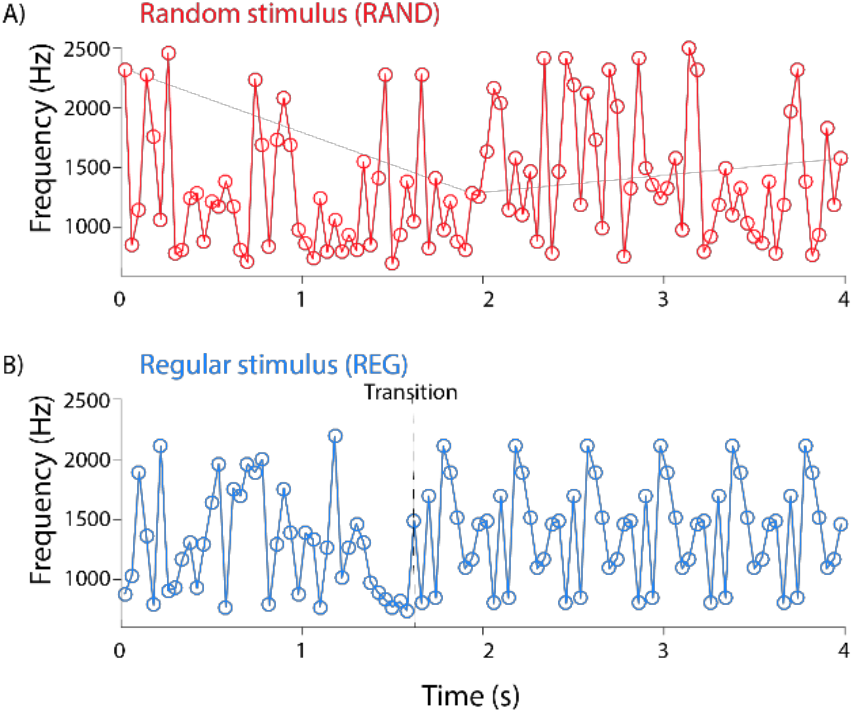
Stimulus conditions and examples of tone presentation in each condition. Examples of auditory stimuli in the Random and Regular conditions are presented in Panel A in red and Panel B in blue, respectively. Each stimulus consisted of ten sets of tones. Each set contained ten tones with a duration of 40 ms/tone. A) For the RAND condition, the tone frequencies for each set were drawn randomly from a pool that ranged from 700 to 2500 Hz. B) For the REG condition, tone frequencies for each set were drawn randomly for the first 4 cycles (1.6 sec). For the remaining sets, 10 tone frequencies were randomly selected and repeated from cycles 5 through 10 (1.6-4 sec). In other words, the REG stimulus transitioned from random to regular, indicated by a vertical dotted line in Panel B.

### 2.4. EEG recording

Participants were instructed to remove any electronic devices, watches or hair clips, before each recording session. EEG signals were recorded at 1024 Hz sampling rate using 16 Ag/Ag-Cl electrodes (Fp1, Fp2, F3, Fz, F4, C3, Cz, C4, T7, T8, P3, Pz, P4, O1, Oz, O2) and left and right mastoid electrodes (Bio-Semi; 208 Hz low-pass filter). Electrodes were referenced to a monopolar reference feedback loop, connecting a driven passive sensor and a common-mode-sense (CMS) active sensor, both located posterior on the scalp. The experiment was conducted in a sound attenuated booth.

### 2.5. EEG data analysis of the sustained response

An elliptic filter was utilized to remove line noise (60 Hz) from the raw data. Offline filtering of the raw data consisted of high-pass filtering at a cutoff of 0.7 Hz (2449 points, Hann window) and low-pass filtering at 22 Hz (211 points, Kaiser window). Data was then down sampled to 512 Hz and re-referenced to the averaged mastoid electrodes. Data was subdivided into five second epochs time-locked to sound onset (−0.8 to 4.8 sec). Independent components analysis (runica method: Makeig, J. Bell., Jung, & Sejnowski, 1996; logistic infomax algorithm: Bell & Sejnowski, 1995; Fieldtrip implementation: Oostenveld, Fries, Maris, & Schoffelen, 2011; RRID:SCR_004849) was employed to identify components related to blinks and horizontal eye movements. The sustained response was investigated using the low-pass filtered data only, meaning that high-pass filtering was omitted in order to retain the low-frequency signal that is reflective of a DC shift and representative of the sustained response (Barascud et al., 2016; Bjorn Herrmann and Johnsrude, 2018; Winkler et al., 2009). However, components from the high-pass filtered data were used to remove activity related to blinks and horizontal eye movements. Epochs that exceeded a signal change of 200 μV in any electrode were excluded from analyses. Offline data analysis was carried out using MATLAB software (8.6; MathWorks, Inc.).

Analyses focused on neural activity, time-locked to the transition to the regular pattern (1.6 s). Single trials were averaged for both conditions (RAND and REG). To baseline correct the response time course, we subtracted the mean amplitude in the pre-stimulus interval (−0.8 to 0 seconds, time-locked to the transition to the regular pattern) from the amplitude at each time point; this was done for each electrode separately. Sustained response data were calculated by averaging signals across the fronto-central-parietal electrode cluster (Fz, F3, F4, Cz, C3, C4, Pz, P3, P4) and calculating the mean amplitude for each condition (RAND and REG) for the 0.8 to 2.4 s time window, that is, the time window in which regularity detection is expected to be completed (Barascud et al., 2016). Selection of the fronto-central-parietal electrode cluster was motivated by previous work on regularity processing showing spatially wide-spread sustained response that is also consistent with the localization of the sustained responses in the auditory cortex and higher-level brain areas.

Next, we calculated the amplitude difference of the sustained response between random and regular (REG-minus-RAND). We calculated the mean amplitude of the sustained response for 4 cycles of regularity, that is cycles 2 to 6 after regularity emergence (0.8 to 2.4 seconds). Previous studies have indicated that the sustained response amplitude is larger (i.e., more negative in EEG) during the REG compared to the RAND conditions (Barascud et al., 2016; Bjorn Herrmann and Johnsrude, 2018; Southwell et al., 2017)

### 2.6. Statistical analysis

We compared the REG-minus-RAND sustained response difference for yCT, aCT, versus PD, in the OFF state, in a one-way analysis of variance (ANOVA) using both frequentist and Bayesian approaches. This was followed by Tukey post-hoc multiple comparisons, to test differences between each of the three groups again using frequentist and Bayesian approaches. Finally, we compared the amplitude of the REG-minus-RAND sustained response difference for each group (yCT, aCT, and PD) against zero, using frequentist and Bayesian one sample t-tests. We next performed an analogous set of analyses on the REG-minus-RAND sustained response difference in the ON Session. Only aCT and PD groups performed regularity detection on dopaminergic therapy. First, we compared aCT and PD in a one-way ANOVA using frequentist and Bayesian approaches. Next, we compared the amplitude of the REG-minus-RAND sustained response difference for each group (aCT and PD) against zero using a) a frequentist one-sample t-test, and b) a Bayesian approach.

It is worth noting that for OFF and ON Bayesian analyses, Bayes factor (BF_10_) ranges are indicative of either support of the null hypothesis (BF_10_ <1) or of the alternative hypothesis (BF_10_ >3), with ranges supporting the strength of either claim (Stefan et al., 2019). Bayesian analysis was performed because it contrasts the probability of the null and the alternative hypotheses in a symmetrical way while giving equal magnitude for either accepting or rejecting the null hypothesis (Dienes, 2014). Bayesian analyses were performed using JASP version 0.8.6.

## 3. Results

### 3.1. Demographic and clinical data

Health and demographic information for both aCT and PD patients are presented in Table 1. The disease duration ranged from 1 to 17 years, with a mean [±standard error about the mean (*SEM*)] duration of 6.78 (± 4.89) years since diagnosis. ACT and PD patients performed similarly on ANART (t34= −0.24; *p* = 0.80), MoCA (*t*_34_= 1.26; *p* = 0.21), and Oxford happiness scale (*t*_34_= 1.34; *p* = 0.18). ACT scored lower than PD patients on the BAI and BDI-II, regardless of the medication state, as has been shown previously (Macdonald et al., 2013; Macdonald and Monchi, 2011; Vo et al., 2014). ACT scored lower than PD patients on our measure of Apathy when they were tested on dopaminergic therapy but there were no group differences in apathy scores in the OFF state. There were no OFF-ON differences for our cognitive/affective measures in either group. Finally, as expected, aCT scored significantly lower than PD patients on the UPDRS Motor sub-scale (*t*_34_ = 20.01 = *p* < 0.001). Performance on UPDRS Motor sub-scale improved with dopaminergic therapy for PD patients (*t*_17_ = 6.67, = *p* < 0.001).

The mean age of our yCT group was 18.6 (*SEM* = 0.5, 8 females). The health, demographic, and cognitive measures described above were not collected for the yCT group.

### 3.2. Sustained response in YCT, ACT, and PD patients OFF medication

Neural sustained response for both conditions are represented for each group (Fig.3). The effect of Group was investigated on the sustained response difference between REG-minus-RAND conditions in the OFF Session in a one-way frequentist ANOVA and a Bayesian ANOVA. We found a highly significant main effect of group using both approaches (*F_2,49_* = 6.533, η_p_^2^ = 0.211, *p* = 0.003; BF_10_ = 13.203). Post-hoc comparisons revealed that the difference REG-minus-RAND (mean±*SEM*) for yCT (−3.457±0.633) was significantly larger than for aCT (−1.066±0.436; *t*_32_ = 3.566; *p* = 0.002; BF_10_ = 11.681). This provided strong support for the difference between yCT and aCT. In contrast, the difference in REG-minus-RAND for yCT and PD groups was only marginally significant or based on the Bayes Factor was considered anecdotal strength evidence of yCT-PD difference (−1.855±0.323; *t*_32_ = 2.389; *p* = 0.053; BF_10_ = 2.448). However, there was no significant difference between the sustained response between aCT and PD patients (*t*_34_ = 1.213; *p* = 0.451; BF_10_ 0.727), with the Bayes Factor supporting the null hypothesis. One sample t-tests on the difference between regular and random (REG-minus-RAND) have revealed a significant difference for yCT, aCT, and PD off medication (Table 2). This was further supported by Bayesian one sample t-tests (Table 2). Significant differences in one sample t-tests indicate that the amplitude difference between REG and RAND conditions is significantly different from zero for all three groups off medication, evidence of the neural response to regularity detection.

**Figure 3:**
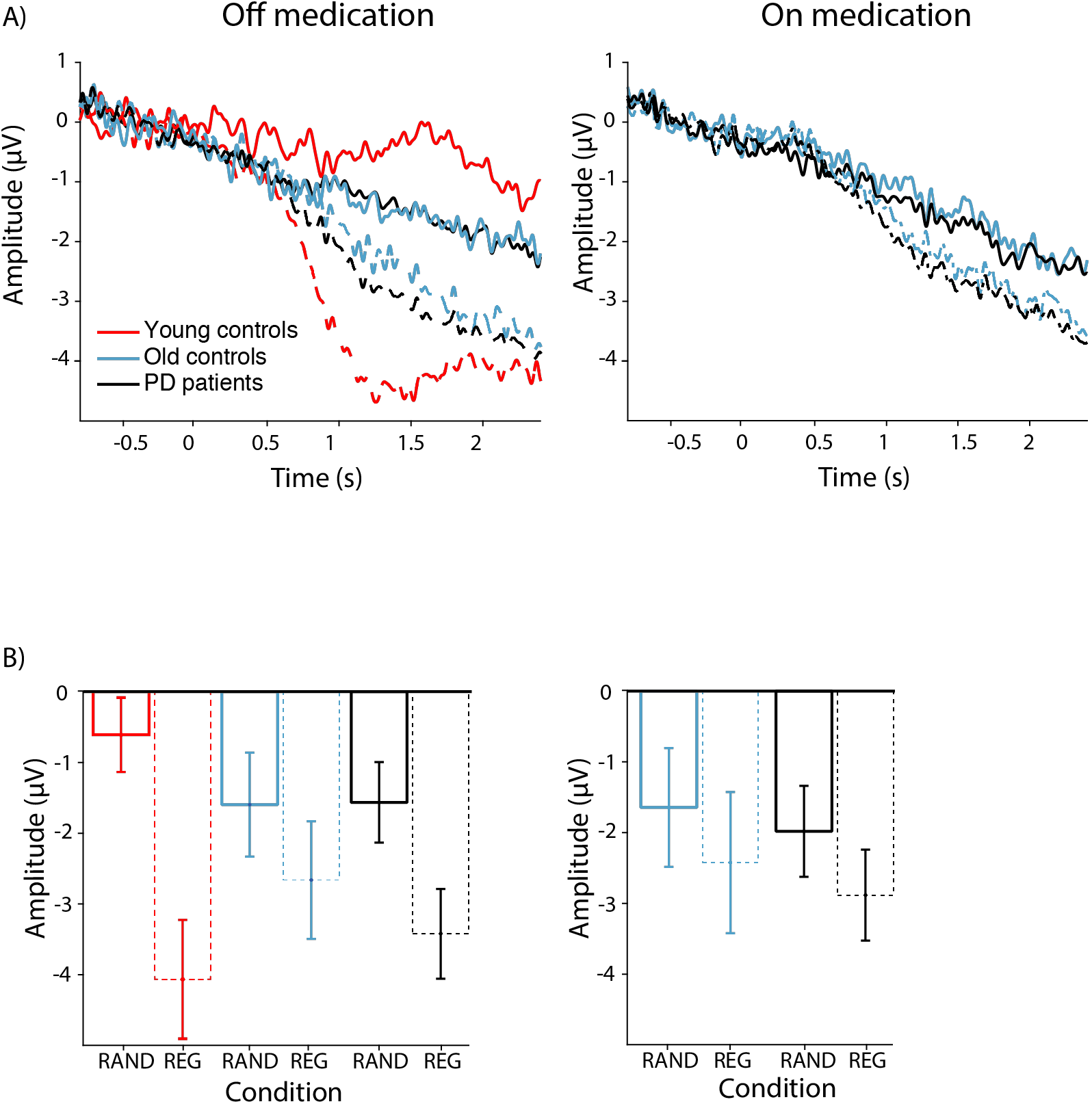
Panel A presents the neural sustained response time course for REG versus RAND from 0.8 seconds prior to the emergence of regularity up to 2.4 seconds following the emergence of regularity in yCT, aCT, and PD in the OFF Session on the Left and in aCT and PD in the ON Session on the Right. REG neural response time courses are presented in dotted lines whereas RAND neural response time courses are presented in solid lines. Panel B presents the mean of the sustained response during the analysis window, which is 0.8 to 2.4 seconds following regularity, for yCT, aCT, and PD in the OFF Session on the Left and for aCT and PD in the ON Session on the Right. The neural sustained response for REG-minus-RAND was attenuated in aCT and PD groups in the OFF Session. The REG-minus-RAND neural sustained response was no longer signficant for aCT and PD groups in the ON Session.

**Figure 4:**
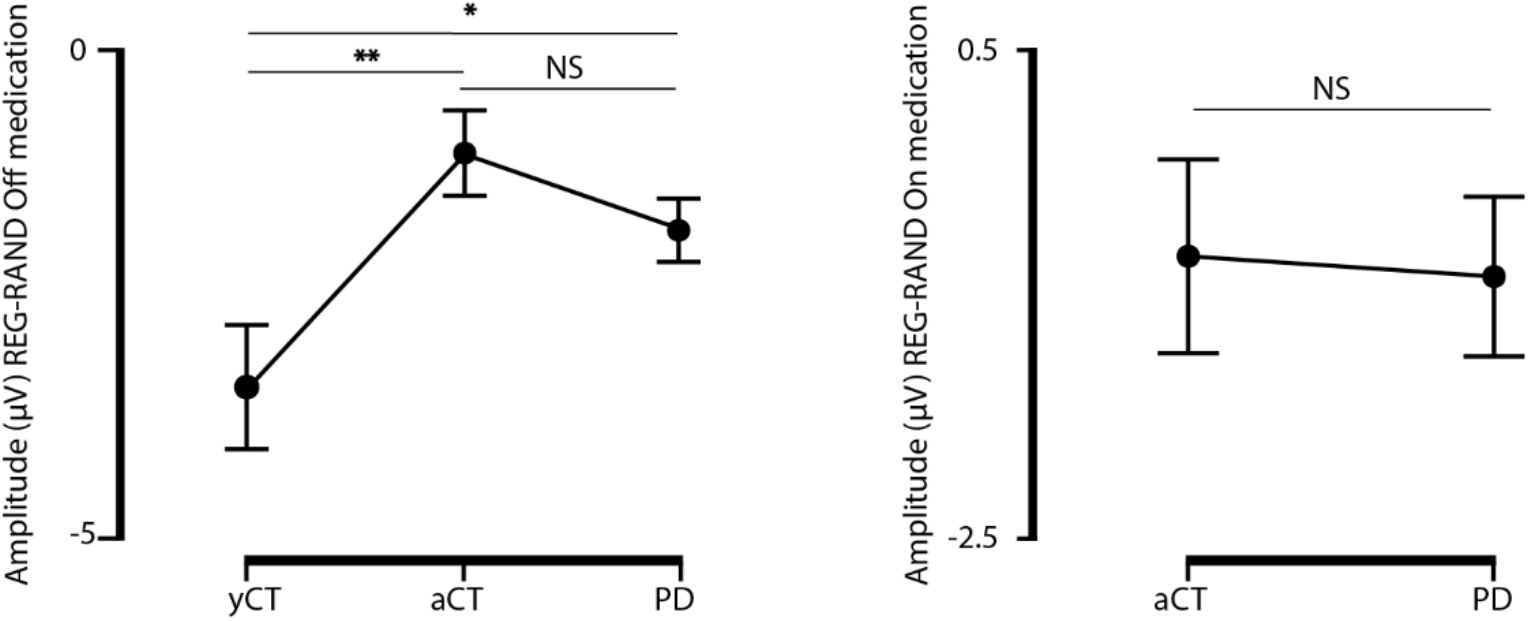
ANOVA on the neural sustained response difference, REG-minus-RAND for yCT, aCT, and PD off dopaminergic medication (Left), and aCT and PD patients on dopaminergic medication (Right) are presented. Off medication (Left), yCT display a larger sustained response (i.e., REG-minus-RAND EEG amplitude difference) compared to aCT and PD patients. On medication (Right), both aCT and PD patients displayed no significant sustained neural response difference for REG-Minus-RAND. yCT = Young healthy Controls, aCT = Aged healthy Controls, PD = patients with Parkinson’s disease. \**p*=*0.05*, ** *p*<*0.01*, NS = Not significant.

**Table 2:** One sample *t*-tests comparing the difference in neural sustained response between REG and RANDfor all groups (yCT, aCT, PD) off and on medication, relative to zero.

|  | One Sample T-Test and Bayesian One Sample T-Test |  |  |  |
| --- | --- | --- | --- | --- |
| | $t$ | $df$ | $p$ | BF<sub>10</sub> |
| yCT off medication * | -5.454 | 15 | < .001 | 408.795 |
| aCT off medication * | -2.444 | 17 | 0.026 | 2.441 |
| PD off medication * | -5.729 | 17 | < .001 | 984.685 |
| aCT on medication | 1.330 | 17 | 0.201 | 0.518 |
| PD on medication | -1.867 | 17 | 0.079 | 1.014 |

Note. Student's $t$ -test.
yCT = Younger controls; aCT = Age matched controls; PD = patients with Parkinson's disease.
df = degrees of freedom; BF<sub>10</sub> = Bayes Factor
\* Significant difference

YCT did not perform the task on L-dopa. For this reason, investigations in the ON state were carried out using data only from aCT and PD patients. One-way frequentist and Bayesian ANOVAs on neural sustained response difference for REG-minus-RAND contrasting aCT and PD revealed no main effect of group (*F_1,34_* = 0.29, η_p_^2^ = 0.001, *p* = 0.873; BF_10_ = 0.325). The Bayes Factor supported the null hypothesis. Further, one sample t-tests did not reach significance with respect to the difference between REG and RAND for both aCT and PD in the ON Session (Table 2). That is, the amplitude difference of REG-minus-RAND for both groups, aCT and PD, in the ON Session were not significantly different from zero. The Bayes Factor provided only anecdotal level evidence in support of the null hypothesis.

## 4. Discussion

In this study, we found that the neural sustained response to the introduction of regularity of auditory signals was present in all groups but was attenuated significantly for aCTs and marginally for PD patients relative to yCTs. There were no differences in the amplitude of the sustained response for REG-minus-RAND between aCTs and PD patients. This suggests that regularity processing worsens with aging but the pathophysiological processes that produce PD do not adversely impact the fundamental function of detecting regularity in sound beyond the effects of aging. Dopaminergic therapy eliminated the significant enhancement in EEG sustained response that occurs for REG-minus-RAND in both PD patients and aCT. These findings suggest that aging, but not PD *per se*, affect the fundamental function of auditory regularity detection. An important finding of this study was that dopaminergic therapy further worsened regularity detection. This occurred in both PD and aCTs.

This is the first study to investigate regularity processing in aging and the first to find a deficit in the neural response related to regularity detection for healthy elderly participants compared to healthy young controls. This finding is in line with a number of results in healthy elderly participants in auditory processing, particularly related to temporal processing and parsing of stimuli. Aging has been found previously to reduce the ability to properly segregate temporally-adjacent auditory stimuli (Aghamolaei et al., 2018; Rimmele et al., 2015; Stothart and Kazanina, 2016). To analyse busy auditory environments, it is necessary to detect the emergence or onset of novel stimuli. Studies investigating auditory scene analysis in older adults have also proposed an impairment in stream segregation, which could be attributed to auditory sensory memory deficits (Rimmele et al., 2012, 2015), selective attention towards stimuli (Aghamolaei et al., 2018; Alain et al., 1996; Christiansen, 2014; Dinces and Sussman, 2017), or improper inhibitory processes of irrelevant sounds (Snyder and Alain, 2007; Stothart and Kazanina, 2016). Basically, it has been suggested that older adults have deficits in sequential grouping and are less able to capitalize on certain regularities for better stream segregation (Rimmele et al., 2012). Getzsmann and Näätänen (2015) found that older adults have deficiencies in stream segregation potentially because they require more time to properly isolate competing auditory inputs when speech in noise was investigated, with the expectation that noise would obscure parts of the conversation, making it harder to follow (Getzmann and Näätänen, 2015). These deficits in aspects of the auditory scene analysis associated with aging seem related to impairments in detecting patterns and segregating them from random or irrelevant auditory stimuli. Despite these previous studies that hint at a problem in regularity processing and detection, ours is the first study to explicitly investigate this and to show that indeed, the neural response related to presentation of a regular pattern embedded in random auditory stimuli was attenuated in healthy elderly participants relative to young controls. Our finding sheds light on these previous auditory processing deficits that have been noted with aging.

No previous studies have invested the effect of PD on regularity processing. PD is a neurodegenerative disorder with prominent motor dysfunction that is related to the loss of dopamine-producing neurons, particularly in the SNc. We found that PD patients and elderly, healthy, matched controls experienced similar attenuation of the sustained response, assessed with EEG, relative to young controls. Regularity processing seems spared by PD pathophysiological processes in that PD did not worsen the neural response related to regularity processing beyond the main effect of aging. Georgiev et al., (2015) have shown that auditory processing is altered in PD patients. PD patients, tested with medication manipulation, were compared to healthy age-matched controls and have displayed deficits in distractor processing in the auditory modality (Georgiev et al., 2015). Moreover, PD patients tested on medication have shown deficiencies in neural entrainment, with an associated deficit between auditory to motor entrainment (te Woerd et al., 2017a, 2017b, 2015). Few studies have systematically sought to disentangle the effects of PD versus dopaminergic therapy on auditory processing through direct contrasts of performance off and on dopaminergic therapy as we have done in this study. Consequently, no clear conclusions regarding PD or dopaminergic therapy on other auditory processes can be drawn.

In contrast to previous studies, we specifically investigated the effect of dopaminergic therapy on regularity processing in both our healthy elderly controls and in PD patients. We found that dopaminergic therapy attenuated the sustained response to the point that there were no significant differences in the EEG responses for sequences in the RAND and REG conditions for both aCT and PD patients. These effects are in line with the dopamine overdose hypothesis (Cools, 2006; Macdonald and Monchi, 2011). This hypothesis proposes that the effect of exogeneous, dopaminergic therapy on functions are determined by the level of endogenous dopamine in the brain regions that underlie them. In PD, SNc is significantly degenerated and the DS, which it supplies, is substantially dopamine depleted. As a consequence, functions mediated by DS reliably improve with dopaminergic therapy (Birkmayer and Hornykiewicz, 1961; Cotzias et al., 1969; Hiebert et al., 2019; Macdonald and Monchi, 2011). In contrast, the VTA is relatively spared in PD, and the brain regions that it innervates, such as the VS and the limbic and prefrontal cortex, are dopamine replete. There is now a large literature showing that in PD, dopaminergic therapy titrated to remediate the DS and address motor symptoms, actually overdoses VTA-innervated brain regions. As a consequence, the functions of VTA-innervated brain regions are worsened in PD by dopaminergic therapy (Cools et al., 2007; Macdonald et al., 2013; Macdonald and Monchi, 2011; Yang et al., 2018). Dopamine overdose effects have been shown repeatedly in healthy young and elderly controls (Coull et al., 2012; van der Schaaf et al., 2013; Vo et al., 2018; Yang et al., 2018), consequently, symmetrical effects of dopaminergic therapy on functions of VTA-innervated brain regions for PD patients and controls are not at all unprecedented. Our results strongly support the contention that regularity processing *depends upon* one or more VTA-innervated brain region. These findings are complementary to findings in the very limited, existing literature that has investigated the neural mechanisms underlying regularity processing. To date, few studies have investigated the brain regions that mediate regularity detection using EEG, MEG, and fMRI (Barascud et al., 2016; Sohoglu and Chait, 2016; Southwell et al., 2017; Teki et al., 2016) in healthy young controls. These studies using neuroimaging found *correlations* between regularity processing and activation in hippocampus and IFG—all VTA-innervated brain regions.

Detection in auditory stimuli of repeating sequences of sounds, or regularities, is a fundamental process that allows us to navigate complex environments. Regularity processing occurs even without attention or consciousness (Chait et al., 2012; Rohenkohl et al., 2011; Southwell et al., 2017). In keeping with this, in our study, participants attended to a subtitled movie without sound, while auditory tones in random or regular arrangements were simultaneously presented. For regular patterns of tones, a sustained enhancement of the neural response, measured with EEG, occurred relative to the presentation of random tones. This sustained response that was observed in the OFF condition for aCT and PD groups was effectively abolished by dopaminergic therapy. Our results, therefore, suggest that dopaminergic therapy interferes with a fundamental process which facilitates the recognition of auditory signals relevant due to their regularity from random, meaningless tones. Learning associations among stimuli (Gotham et al., 1988), responses, and rewards (Cools, 2006; Gotham et al., 1984; Hiebert et al., 2019; Kish et al., 1988; MacDonald et al., 2013; Swainson et al., 2000; Yang et al., 2016), as well as sequences and cognitive processing speed (Poewe et al., 1991) are the most common cognitive functions that are worsened by dopaminergic therapy. Detection of patterns as well as repeating stimuli or events seem prerequisite for the aforementioned forms of learning. This raises the possibility that learning deficits induced by dopaminergic therapy in PD could be related to fundamental impairment in regularity processing and detection. Indeed, noting and exploiting statistical regularity in the environment is an element of many cognitive functions (learning, memory, and intelligence).

## 5. Conclusion

Regularity processing is a fundamental function in auditory scene analysis and negotiation of complex auditory environments. Deficits in detecting regular patterns can instigate a number of cognitive impairments including most forms of association and sequence learning as well as language processing and multi-sensory integration. We found that aging impaired the neural signal associated with regularity detection assessed with EEG. PD did not worsen regularity processing above the main effect of age, given that PD is an aging-related disorder. Beyond these deficits, however, dopaminergic therapy (i.e., regular regimens for PD patients; a single dose of L-dopa for controls) abolished the sustained response in PD patients and healthy elderly participants such that the EEG responses for regular and random sequences of tones were no longer significantly different. This pattern of deterioration of function related to dopaminergic therapy is a well-documented phenomenon attributed to dopamine overdose of dopamine-replete brain regions due to their innervation by the VTA which is spared in PD and normal aging. This coheres with the limited literature on regularity processing, relating this fundamental process to a number of VTA-innervated brain regions (e.g., hippocampus and IFG). Most important, the elimination of the neural response associated with regularity processing could instigate a number of cognitive impairments such as association learning and speech processing, that depend on regularity processing. This understanding should be weighed against motor symptoms in titrating dopaminergic medication in PD patients.

## Acknowledgment

This study was supported by a Canada Research Chair (CRC) Tier 2 in Cognitive Neuroscience and Neuroimaging to P. A. MacDonald, an Opportunity Grant from the Academic Medical Organization of Southwestern Ontario to P. A. MacDonald, an Internal Research Fund Grant from Lawson Research Institute to P. A. MacDonald. BH was supported by a BrainsCAN postdoctoral fellowship (Canada First Research Excellence Fund; CFREF). We would also like to thank Nole Hiebert and Josh Hoddinott for reviewing this manuscript.

